# Integrated profiling of single cell epigenomic and transcriptomic landscape of Parkinson’s disease mouse brain

**DOI:** 10.1101/2020.02.04.933259

**Authors:** Jixing Zhong, Gen Tang, Jiacheng Zhu, Xin Qiu, Weiying Wu, Ge Li, Xiumei Lin, Langchao Liang, Chaochao Chai, Yuying Zeng, Feiyue Wang, Lihua Luo, Jiankang Li, Fang Chen, Zhen Huang, Xun Xu, Shengping Tang, Shida Zhu, Dongsheng Chen

**Author notes:** These authors contributed equally to this work (Jixing Zhong, Gen Tang, Jiacheng Zhu, Xin Qiu, Weiying Wu, Ge Li). Correspondence should be addressed to DSC, SDZ or SPT.

## Abstract

Parkinson’s disease (PD) is a neurodegenerative disease leading to the impairment of execution of movement. PD pathogenesis has been largely investigated, but either restricted in bulk level or at certain cell types, which failed to capture cellular heterogeneity and intrinsic interplays among distinct cell types. To overcome this, we applied single-nucleus RNA-seq and single cell ATAC-seq on cerebellum, midbrain and striatum of PD mouse and matched control. With 74,493 cells in total, we comprehensively depicted the dysfunctions under PD pathology covering proteostasis, neuroinflammation, calcium homeostasis and extracellular neurotransmitter homeostasis. Besides, by multi-omics approach, we identified putative biomarkers for early stage of PD, based on the relationships between transcriptomic and epigenetic profiles. We located certain cell types that primarily contribute to PD early pathology, narrowing the gap between genotypes and phenotypes. Taken together, our study provides a valuable resource to dissect the molecular mechanism of PD pathogenesis at single cell level, which could facilitate the development of novel methods regarding diagnosis, monitoring and practical therapies against PD at early stage.

## Introduction

Parkinson’s disease (PD), known as the second-most prevalent neurodegenerative disease in the world, is predominantly characterized by motor disorders. Patients with PD also show non-motor symptoms including cognition impairments, autonomic dysfunction, hyposmia and so on^1^. PD is presently incurable and as PD gradually progress, the symptoms will eventually deteriorate into severe disabilities. The genetic mechanisms behind have been widely and profoundly studied in the last two hundred years. Accumulating evidences attribute PD pathologies to the malfunction of nigrostriatal dopamine pathway^2^ where dopaminergic neuron in substantia nigra (SN) release dopamine from axon terminals that synapse onto the medium spiny neurons in dorsal striatum. In PD, accumulation of aggregated *α*-synuclein (*α*-syn) leads to progressive degeneration of dopaminergic neurons in nigrostriatal dopamine pathway^3^, resulting in the remarkable reduction of dopamine level and symptomatic motor deficits in PD. Plenty of studies have made great progress in disentangling the link between *α*-syn oligomerization and neuron death, revealing pathways including proteostasis, mitochondrial dysfunction, neuroinflammation and so on^4^. Nevertheless, the majority of previously studies were conducted at tissue level. Wassouf’s team performed RNA-seq on 6-month-old mice overexpressing *SNCA* (*α*-syn encoding gene) and found that striatal gene expression profiles were greatly disrupted, whose functions were primarily related to neuroinflammation and synaptic plasticity^5^. Richard et. al. combined bulk and single-cell RNA-seq for iPSC-derived dopamine neurons with a *GBA* mutation and identified HDAC4 as a potential therapeutic PD target^6^. Given the fact that intricate interplay across and within cell types jointly contributes to PD pathogenesis^7^, it is essential to interrogate cell-resolution information in order to unveil the possible PD-related and PD-causing molecular circuits. Single cell sequencing technologies have facilitated the researches in various fields, providing us an opportunity to capture subtler changes that may be masked by bulk sequencing.

Of note, due to synaptic plasticity, 80% of dopaminergic neurons in SN have been lost before any diagnosable symptoms of PD occur^8^. Hence, this raises the necessity to dissect cellular changes in the early stage of PD and identify corresponding markers, ultimately allowing the development of novel early interventions. Nigrostriatal dopamine pathway malfunction attributes largely to PD pathologies^9^. Besides, cerebellum is the pivot of motor coordination but has been often overlooked in Parkinsonism. Therefore, with the aim to explore the deregulation in early stage of PD transcriptionally and epigenetically, we applied single-nucleus RNA sequencing (snRNA-seq) and sci-ATAC-seq on 6-month old PDm and WTm brain regions including midbrain striatum and cerebellum. We revealed the extensive, yet region-specific dysfunctions regarding proteostasis, channelosome, extracellular environment and neuroinflammation. Specifically, we noticed that the capability of releasing neurotransmitters of dopaminergic neuron may be impaired. We also discovered the transcriptional alterations of dopaminergic neurons showed great divergences compared to that of late stage PD, highlighting the significance of our datasets. Besides, through combining transcriptomic and epigenetic profiles, we proposed three classes of biomarkers for PD that may be advance the diagnosis, monitoring and therapies development.

## Result

### Single-nucleus profiling of cerebellum, midbrain and striatum transcriptionally and epigenetically

Cerebellum, midbrain and striatum sample were dissected from 6-month old *α*-synuclein A53T transgenic mouse (PDm) and matched wildtype mouse (WTm) to perform the single-nucleus sequencing (Figure 1a). Generally, we obtained 46,174 individual transcriptomic profiles, 46,146 of which were retained after filtering. Cells from cerebellum, midbrain and striatum were relatively even: 13,774 cerebellar cells, 14,117 midbrain cells and 18,255 striatal cells (Figure S1a). Also, we applied sci-ATAC-seq to investigate the whole chromatin accessibility of three brain regions mentioned above. Consequently 28,347 cells (4,914 from cerebellum, 11,224 from midbrain and 12,209 from striatum) passed stringent filtering standards, resulting in the median of unique fragments for cerebellum (Figure S1f), midbrain and striatum were 2,201, 4,959 and 5,651 respectively (Figure S1f).

**Figure 1.**
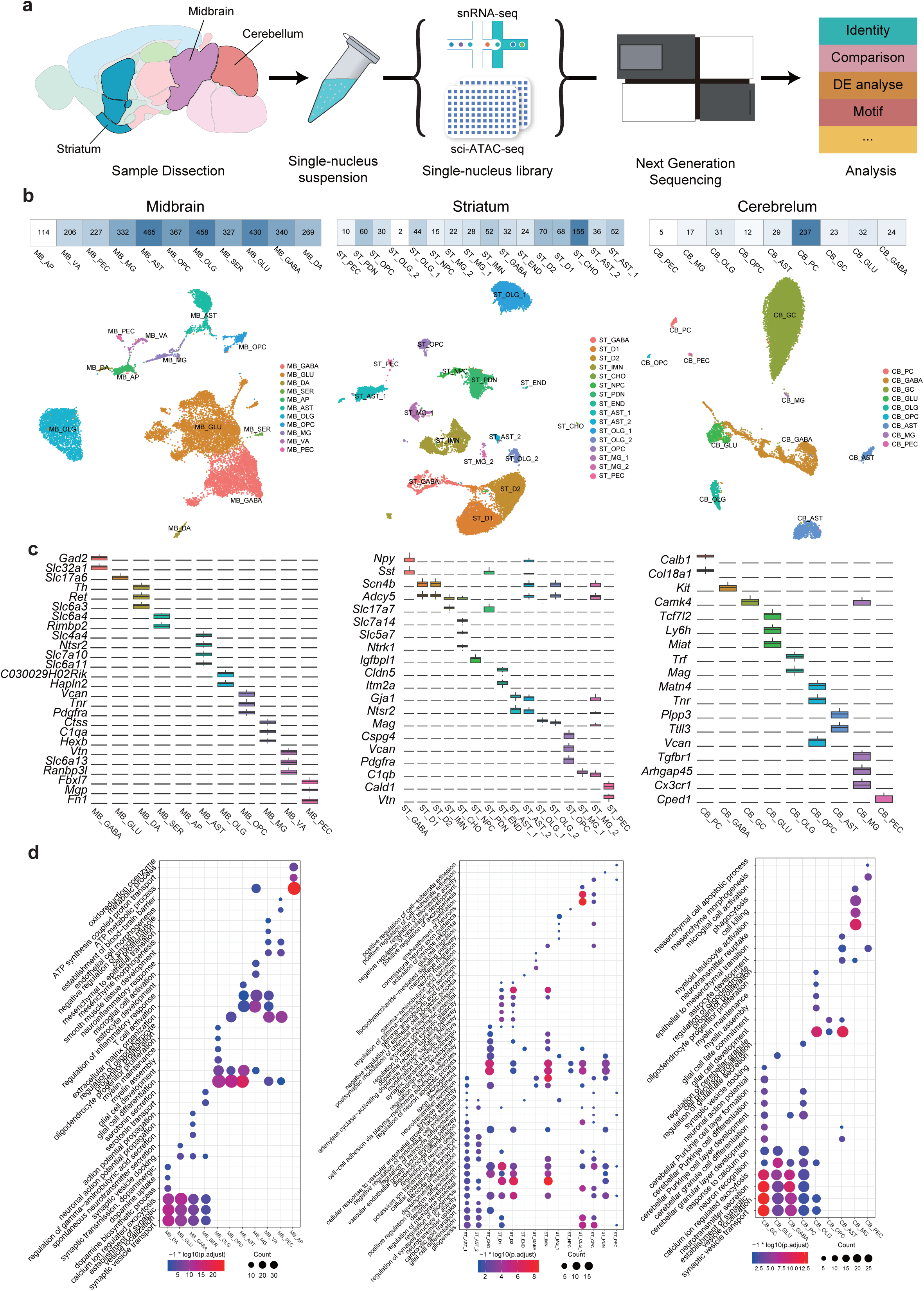
a. Scheme of the experimental and bioinformatic design in our study. b. Unsupervised clustering of snRNA-seq datasets of three brain regions with cells colored by cell types. Left: midbrain; middle: striatum; right: cerebellum. Heatmap on top shows the number of cell type-specific DEGs. c. Boxplots showing the expression patterns of cell type-specific DEGs in midbrain (left), striatum (middle) and cerebellum (right). d. Selected GO terms enriched in each cell type. Dot size indicates the number of genes enriched in certain term while color indicates *P*-value. Left: midbrain; middle: striatum; right: cerebellum.

### Heterogeneity of gene expression in PDm and WTm brain revealed by snRNA-seq

To classify major cell types, we combined cells from both PDm (8,921 cells) and WTm (5,206 cells), followed by unsupervised clustering. Based on reported markers, we assigned them to 10 cell types in midbrain including GABAergic neuron (MB_GABA), dopaminergic neuron (MB_DA), glutamatergic neuron (MB_GLU), serotonergic neuron (MB_SER), oligodendrocyte (MB_OLG), oligodendrocyte precursor cell (MB_OPC), astrocyte (MB_AST), microglia (MB_MG), vascular cell and pericyte (MB_PEC) (Figure 1b). MB_GABA specifically expressed *Slc32a1*, *Gad1* and *Gad2* while MB_GLU showed enriched expression of *Slc17a6* and *Slc17a7*. MB_DA were identified according to canonical markers *Th* and *Slc6a3*. We also identified MB_SER that was consisted of a relatively small portion of cell, by distinct expression of *Slc6a4*. MB_OLG and MB_OPC shared oligodendrocyte features by expressing *Olig1*, *Cldn11* and *Mbp*, on top of which MB_OPC specifically expressed *Cspg4* and *Pdgfra*. MB_ASTs exhibited high expression of astrocytic markers, including *Aqp4*, *Mfge8* and *Fgfr3*. MB_MG distinguishably expressed *Ctss*, *Ptprc*, *Laptm5* etc. MB_PECs and vascular cell were characterized by the expression of *Rgs5* and *Lum*, respectively. Likewise in striatum, we analyzed 18,255 striatal cells (5,444 from WTm and 12,811 from PDm) and successfully captured major striatal cell types: medium spiny neuron (ST_MSN_D1, ST_MSN_D2), cholinergic interneurons (ST_CHO), GABAergic interneuron (ST_GABA), immature neuron (ST_IMN), neural progenitor cell (ST_NPC), oligodendrocyte (ST_OLG_1, ST_OLG_2), oligodendrocyte precursor cell (ST_OPC), astrocyte (ST_AST_1, ST_AST_2), microglia (ST_MG_1, ST_MG_2), endothelial cell (ST_END) and pericyte (ST_PEC).

Specifically, striatal MSNs can be further divided into two known categories based on the expression of D1-type (*Drd1*) and D2-type receptors (*Drd2*). We identified D1-type MSNs and D2-type MSNs highly expressing *Drd1*, *Tac1* and *Drd2*, *Penk* respectively. Interestingly we noticed small subpopulations of AST (ST_AST_2), OLG (ST_OLG_2) and MG (ST_MG_2) expressing either D1 or D2-type receptors and were adjacent to MSNs in the UMAP plot. We inferred that ST_AST_2, ST_OLG_2 and ST_MG_2 may surround MSNs spatially under physiological condition and dopamine receptors were induced by the consistent dopamine stimulation to coordinate proper dopamine signaling. Supporting our results, AST has been reported to express dopamine receptors and transporters, through which dopamine can signal on and trigger complex downstream intracellular cascades^10^. ST_GABA can be further classified into known subpopulations, including *Npy^+^Sst^+^* (C2_SOM-1, C4_SOM-2, C8_SOM-3), *Vip^+^* (C6_Vip), *Pvalb^+^* (C1_Pvalb), *Th^+^* (C7_Th) and *Cck^+^* (C5_Cck) (Figure S3g, h). Aside from that, we noticed C0 and C3 highly expressing *Foxp2* and *Igfbp4*, which may be responsible for specific functions in striatum (Figure S3g, h). ST_IMNs were characterized by distinct expression of markers for immature neurons, such as *Dcx*, *Tbr1* and *Neurod1*. ST_CHOs expressed *Chat, Ache* and *Slc18a3*. ST_ENDs specifically expressing *Ly6c1*, were considered as endothelial cells (Figure S2b).

In terms of cerebellum, we generated 13,774 cells (3,452 from WTm and 10,322 from PDm), covering a comprehensive range of cell types comprising Purkinje cell (CB_PC), GABAergic interneuron (CB_GABA), granule cell (CB_GC), glutamatergic neuron (CB_GLU), oligodendrocyte (CB_OLG), oligodendrocyte precursor cell (CB_OPC), astrocyte (CB_AST), microglia (CB_MG) and pericyte (CB_PEC) (Figure S2c). CB_PCs showed recognizable expression of *Calb1* and *Car8*, both classic markers for Purkinje cells. Although CB_GCs and CB_GLUs were both glutamatergic neurons, the former specifically expressed *Pde1c* and *Slc17a7*, whereas the latter expressed *Slc17a6*, *Meis2* and *Lhx9*. Particularly in CB_AST, Bergmann glia cells were identified with high expression of *Gdf10*, apart from classic AST marker *Slc1a3* and *Aqp4* (Figure S4e, f).

We next inspected the expression of cell type differentially expressed genes (DEGs) and observed distinct transcriptional patterns across cell types. We noticed that many reported cell type markers were also identified as cell type DEGs, aside from which we also observed other DEGs that exhibited distinct expression patterns (Figure 1c). In midbrain, MB_DA specifically expressed *Ret* and MB_SER showed distinct expression of *Rimbp2*. CB_PCs highly distinguishable expression of *Col18a1* that has been previously reported to be essential for the guidance of climbing fiber terminals onto PC^11^. ST_MSNs specifically expressed *Scn4b* and *Adcy5*. Functional enrichment analysis of genes differentially expressed by each cell type provided further evidences for cell type assignment (Figure 1d). For example, MB_GABA, MB_GLU, MB_DA and MB_SER-specific genes were enriched in “synaptic vesicle transport”, “establishment of synaptic vesicle localization” and “calcium ion regulated exocytosis”, all of which indicated mature neuron functions. Specifically, MB_DA and MB_SER had GO terms “dopamine uptake” and “serotonin secretion” respectively, matching the pre-defined cell identities. MB_OLG-specific genes have “myelin assembly” and “myelin maintenance” related processes, whereas MB_OPC-specific genes were distinctly enriched for terms like “extracellular matrix organization”. MB_AST-specific genes were closely associated with “astrocyte development” and neuroinflammatory pathways, which were consistent with AST functionalities. MB_MG governs the environmental homeostasis in brain, with DEGs enriched with GO terms “microglial cell activation” and “neuroinflammatory response”. Apoptosis showed significant enrichment of ATP metabolic processes. Additionally, MB_PEC and vascular cells were associated with the “establishment of blood-brain barrier” and “mesenchymal to epithelial transition”. In cerebellum, we noticed “cerebellar granular layer development” were associated with CB_GC and CB_AST, indicating granular layer development not only account for CB_GC, but also CB_AST.

### Alterations of gene expression in Parkinson’s disease mouse brain

We next interrogated the molecular changes in different brain regions at transcriptome level by pair-wise comparison of PDm and WTm within same cell type. A large scale of transcriptional alterations was quantified between PDm and WTm, indicating the occurrence of PD dysregulation. In total, 2,116 up-regulated and 469 down-regulated DEGs were identified in midbrain (Figure 2b). We inferred from the number of DEGs that all major cell types were affected by PD pathology. Among those down-regulated DEGs, we found *Pbx1*, *Mdm4* and *Clk1* that have been reported to associate with dopamine neurons dysregulation in PD. *Pbx1* plays a vital part in midbrain dopaminergic neuron development which has been reported to be impaired in PD^12^. *Mdm4* has the capability to restrict the transcriptional activity of tumor suppressor p53^13^ and p53 inhibitors are highly effective in protecting midbrain dopamine neurons and preserving motor function in PD mouse model^14^. Besides, alternative splicing dysregulation of *Mdm4* can lead to the death of motor neurons^15^, which is one of the most prominent pathological features of PD. Additionally, *Clk1* deficiency inhibited autophagy in dopaminergic neurons both *in vitro* and *in vivo* via regulating intracellular autophagy-lysosome pathway^16^. Consequently, our observations of *Pbx1*, *Mdm4* and *Clk1* may point to their essential but previously elusive roles in early stage of PD. On the other hand, the numbers of DEGs in other neuronal types, both MB_GABA and MB_GLU included, remarkably outnumbered that of MB_DA. As the profiled MB_GABA and MB_GLU mainly reside outside SN, neurons perturbations emerge in other subregions within midbrain prior to MB_DA were substantially affected and may even contribute to the later malfunction in SN. In striatum, prominent number of DEGs were identified in ST_MSN (699 in ST_MSN_D1 and 676 in ST_MSN_D2) (Figure S3b). Dorsal striatal MSN receives dopamine from midbrain DA and signal through direct and indirect pathway to coordinate body movement. Thus, MSNs has been transcriptionally affected preceding DA degeneration, possibly by the shrinkage of dopamine level. In cerebellum, we also inspected gene-level alterations by the identification of DEGs. In total, we identified 1,636 upregulated and 233 downregulated DEGs in cerebellum (Figure S4b), most of which were specific to one cell types (Figure S4g).

**Figure 2.**
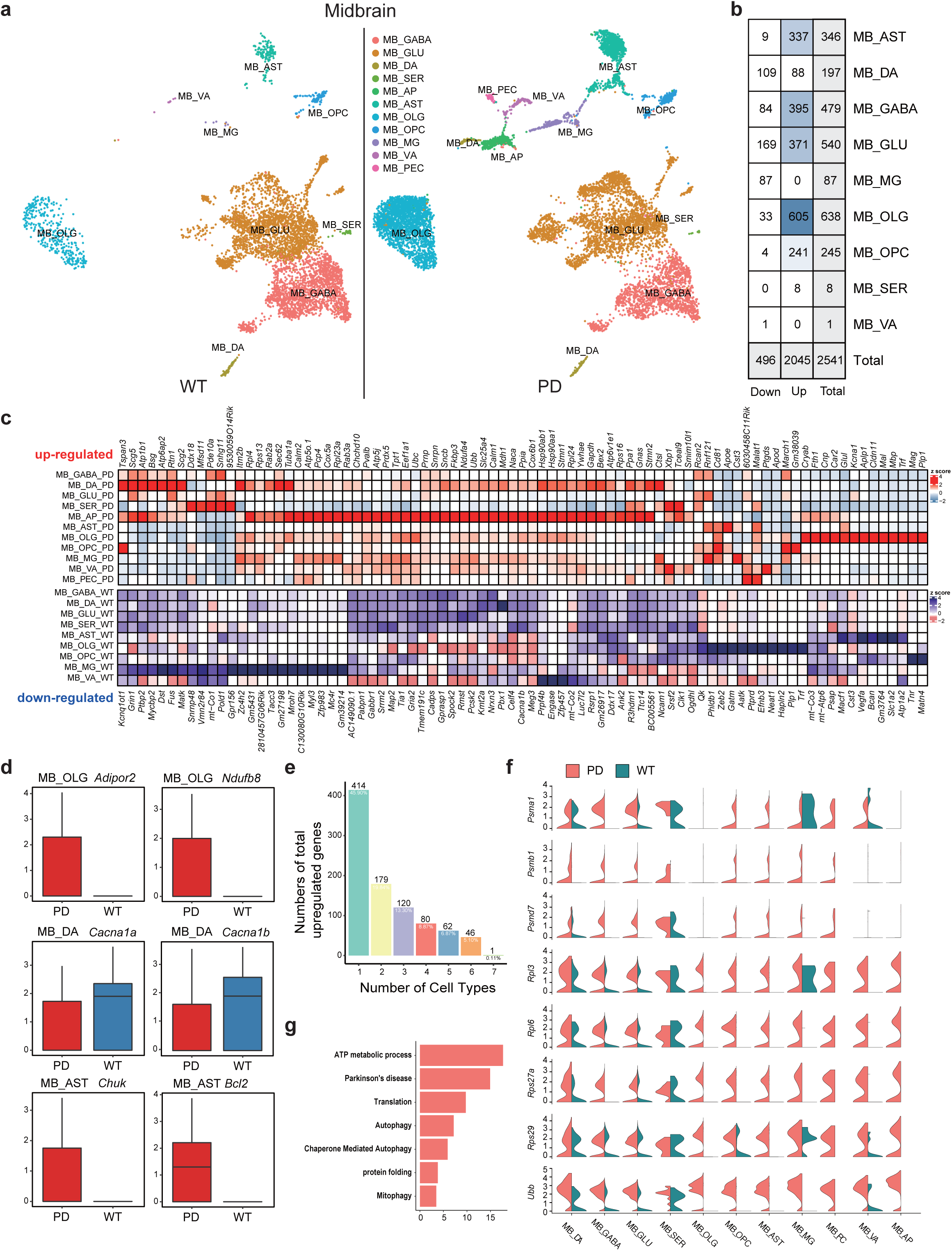
a. Unsupervised clustering of snRNA-seq datasets of midbrain, with PD dataset shown in left panel WT dataset shown in the right. Cells are colored by cell type. b. The number of cell type-specific genes c. Expression profiles of cell type-specific genes. d. Expression profiles of selected DEGs e. The numbers of total upregulated genes (y axis) as a function of the total number of cell types in which the upregulation occurs. f. Selected GO terms enriched in genes with over three times of occurrence across different cell types. g. Expression profiles of autophagy-related genes.

### Perturbation of protein homeostasis network in PDm brain

In midbrain, we noticed that a remarkable portion of PD up-regulated DEGs were shared across cell types (Figure 2e), indicating some biological processes were disrupted commonly in different cell types. Given the machinery that genes execute biological functions collectively as organic modules, we employed functional enrichment analysis to explore the features of genes that shared by greater than three cell types^17^. Dysregulated pathways were highly overlapped between midbrain and striatum and the overlapped events were collectively involved in cellular proteostasis process. Active ATP metabolism and upregulated ribosomal protein encoding genes (RPL and RPS genes) suggested elevated activities in protein synthesis and may linked to neurotoxic aggregation. Additionally, increased expressions of protein degradation related genes were observed, including through ubiquitin-proteasome system and autophagy-lysosome system. This was supported by the extensive upregulation of heat shot protein genes, proteasomal genes and ubiquitin genes (*Psma1*, *Psmb1, Psmd7*, *Rpl3*, *Rpl6*, *Rps27a*, *Rps29*, *Ubb*) (Figure 2f). Surprisingly, *Cryab*, which has been reported as an inhibitor of autophagy activities and an anti-apoptotic factor whose malfunction may accelerate the deterioration of neuroinflammation and demyelination^18^, showed preferential expression in PDm^19^, indicating the underlying regulatory mechanism of protein degradation is more intrinsic than expected.

In PD down-regulated DEGs within midbrain, DEGs are more cell type-specific, with about 70% of total down-regulated DEGs perturbed in single cell type. We performed functional enrichment analysis and found that DEGs in glia cells didn’t converge to certain biological functions. While in neuronal cell types, by assessing the functionalities of DEGs, we found that mRNA processing, especially mRNA splicing events may be disrupted in PD (Table S2). Relevant genes included (but not limited to) genes that encode RNA-binding protein (*Rbm25*), spliceosome (*Srsf2*) and RNA helicase (*Ddx17*). Aside from that, CELF family were observed to downregulated be at mRNA level. CELF protein family has long been associated with alternative splice sites selection and were ascribed to the pathologies of multiple neurological disorders, including some neurodegenerative diseases^20–22^. Keeping with our findings, a differential co-expression analysis research on SN tissue, an alternative isoform of *SNCA* with long 3’UTR is preferentially linked to Parkinsonism^23^. We assumed that the deregulation of mRNA splicing may be attributed to the alteration in splice sites caused by mutations in *α*-syn encoding sequence. Neurons respond to abnormal *α*-syn aggregates by affected splicing events, may leading to an enhanced resistance to degradation system. Besides, synapse assembly and axonogenesis were suggested to be down-regulated in MB_GABA and MB_GLU.

Specifically, we found that significant disruptions in Rab GTPases family (Table S2), which is engaged in multiple steps of membrane trafficking. For example, *Rab2a* is involved in retrograde trafficking, recycling particles from Golgi back to the endoplasmic reticulum (ER). This retrieval is part of the organellar homeostasis pathway to prevent misfolded proteins from entering Golgi apparatus. Thus, the upregulation of *Rab2a* in PDm, which may promote retrograde trafficking machinery, may be the stress response for *α*-syn aggregation. *Rab3a*, localized to presynaptic termini, is responsible for the tethering and fusion of neurotransmitter vesicles preceding release. It has been reported that RAB3A can interact with mutant *α*-syn but not wild-type *α*-syn^24^, based on which we presumed that the upregulation of *Rab3a* in PDm implied a compensatory effect of the binding of Rab3a and aberrant *α*-syn. We also raised the possibility that this pathological binding can result in diminished release of neurotransmitters. Accordant with our findings, overexpression of *Rab3a* can ameliorate α-syn toxicity in yeast^25^. On top of that, we also observed *Rab26os* was downregulated in neurons (MB_DA, MB_GABA, MB_GLU). *Rab26os* is a lncRNA that encoded by the opposite strand of *Rab26* which holds a pivotal role in excessive vesicles degradation through autophagy. *Rab26os* has not been associated with specific functions yet. Nonetheless, as postulated by the theory that antisense lncRNA may have a role in the regulation of sense gene expression^26^, we proposed the possible existence of the Rab26-Rab26os regulatory network. Therefore, the abated expression of Rab26os may interfere the network homeostasis, regulating the formation of autophagosomes. Of note, cell type-specific divergencies were observed. In MB_AST, *Rab6a* and *Rab6b* were upregulated in PDm, indicating not only *Rab2a* mentioned above, Rab6 complex is suggested to be involved in COPI-independent retrograde trafficking in MB_AST. Besides, *Rab5b* and *Rab7*, which jointly involved in endosome maturation, showed hyperactivated expression in MB_OLG from PDm. Thus, elevated expression of *Rab5b* and *Rab7* may indicate the internalized molecules undergo a quick Rab5 to Rab7 conversion before taking any effects. In line with our results, a recent study observed enhanced expression of Rab5 in PD mouse model overexpressing wild-type *α*-syn^27^.

We also observed similar perturbations in striatum. In summary, beyond the consistence in proportional discrepancies between PDm and WTm, striatum functionally responds to *α*-syn aggregation in a similar fashion to midbrain. Protein metabolism were disorganized with augmented protein synthesis and impaired degradation, worsening the deposition of mutant *α*-syn. The release of neurotransmitters began to be hindered due to the chaos in membrane trafficking network.

### Disruption of ion channel in PDm brain

Ion channels are essential for the realization of neuronal functions by delicately regulating membrane potential. Hence, we wondered whether and to what extent, ion channels are disturbed and contribute to early PD pathogenesis. Intriguingly, *Cacna1a* and *Cacna1b*, promoting neurotransmitter release by inducing Q/P-type and N-type calcium currents respectively^28^, were down-regulated in MB_DA (Figure 2d), implying a defectiveness of MB_DA to release dopamine on downstream targets. It is previously acknowledge that the loss of dopaminergic neurons leads to Parkinsonism, here we proposed that the release of dopamine was also hindered in MB_DAs before the degeneration of DA, thus being culpable for the diminished dopamine level in striatum, particularly at early stage of PD. In ST_MSN from PDm, we observed that *Scn4b*, encoding a subunit of voltage-gated sodium channels that modulate the influx of Na^+^, showed reduced expression in ST_MSNs from PDm (Figure S3f). Thus, we presumed that a diminished cellular Na^+^ concentration in PD ST_MSNs, hindering the occurrence of action potential. Additionally, PD ST_MSNs expressed *Ryr3* at significantly lower level (Figure S3f), which are responsible for the release of Ca^+^ from endoplasmic reticulum into cytoplasm^29^, indicating an aberrant Ca^+^ signal in PD ST_MSNs. Taken together, ST_MSNs in PD condition may be under a relatively inactive state, as reflected by possible obstacles in the formation of action potential and dysfunctional Ca^+^ signal. Specifically, we noticed that *Kcnc3* and *Itpr1* were both down-regulated in CB_PC in PDm (Figure S4h). *Kcnc3* encodes a fast-activating/deactivating potassium voltage-gated channel Kv3.3, which drive the rapid repolarization phase of action potentials while *Itpr1* encodes an intracellular IP3-gated-calcium-release channel and mediates calcium release from the endoplasmic reticulum. The decreased expression of *Kcnc3* would prolong the duration of action potential spikes that might increase cellular calcium influx^30^. Besides, reduced expression of *Itpr1* has been demonstrated to cause aberrant intracellular calcium signaling in Purkinje cells. Collectively, we suspected that cellular calcium level and Ca^+^ dependent signaling may play a crucial role in the development of PD symptoms. In line with our results, Mice lacking *Kcnc3* showed moto disorders such as motor incoordination, muscle twitches and constitutive hyperactivity^31^ and *Kcnc3*-null cerebellar Purkinje cells were recorded perturbed complex spikes^32^. Decreased expression of *Itpr1* has been demonstrated to cause aberrant intracellular calcium signaling in Purkinje cells and mutation in *Itpr1* is associated with pathogenesis of spinocerebellar ataxia ^33^.

### Dysregulation of glutamatergic receptors and transporters in PDm brain

Astrocytes performs diverse functions in central nerve system, such as synchronizing axon activity, maintaining energy metabolism and homeostasis, regulating the extra-neuronal environment *etc*^34^. AST serves as neuroprotector and neurodegenerator dependent on the molecules it release into and take up from extracellular space^35^. Recent insights proposed that astrocytes appear to initiate and/or drive the progress of PD. Observational and experimental studies have indicated that *α*-syn can be taken up by AST and spread to neurons in a non-cell-autonomous manner^7^. According to our dataset, 346 genes were upregulated in PD MB_AST, among which we observed glutamatergic transporters and receptors (*Gnao1, Gria1, Gria2, Plcb1, Slc1a2, Slc38a2, Dlgap1*) (Table S2). Glutamatergic transporters serve to keep low glutamate level in ambient extracellular environment while upregulation of astrocytic glutamatergic receptors have been reported to be capable of causing calcium-dependent release of gliotransmitters^36^. In line with this, we also found a regulator of cytosolic calcium concertation, *Saraf*, being upregulated in PDm. Likewise in striatum, we observed two major glutamate transporters, GLAST and GLT-1, encoding by *Slc1a3* and *Slc1a2* respectively, displayed diminished expression in ST_AST_1 from PDm (Figure S3f), implying an impediment of glutamate uptake by PD ST_AST_1 which may lead to increased extracellular glutamate concentration and overexcitation of surrounding neurons. In addition, we observed augmented expression of *App* and *Apoe* in ST_AST_1 (Table S2), which may result in Aβ formation and deposition that known to affect the activity of GLAST and GLT-1 and downregulates glutamate uptake capacity of astrocytes^37^. Thus, we speculated that the ability to reuptake extracellular glutamine of AST were impaired in early PD striatum, possibly by misfolded protein deposition.

### Increased activity of NFκB signaling pathway in PDm brain

Neuroinflammation is a salient signature of PD neuropathology. Evidences from previous observations indicate α-syn aggregation induces immune responses in PD patients^38, 39^ and neuroinflammatory activities can facilitate α-syn misfolding^40^, forming a self-aggravating loop. The deposition of *α*-syn causes AST to produce pro-inflammatory cytokines and activate MG, suggesting a plausible role of AST in the initiation of PD^23, 24^. Given that AST and MG are two primary cell types that closely involved in neuroinflammation^43^, we next focused on AST and MG to explore the neuroinflammatory reactions at early stage of PD. Transcriptomic changes of MB_MG under pathological state were not compelling, with only 87 down-regulated genes and none up-regulated genes (Figure 2b). We also noticed that activated MG markers *Aif1* and *Slc2a5* showed overt enriched expression in MB_MG from PDm, based on which we inferred that MB_MG was activated and under the state of proliferation. On the other hand in MB_AST, we noticed increased activities in NFκB signaling pathway due to the upregulation of *Bcl2*, *Csnk2a1* and *Chuk* in PDm (Figure 2d). *Chuk* encodes IKKα whose activation is essential for the activation of NFκB canonical pathway by phosphorylating the IκBα protein^44^. *Bcl2* and *Csnk2a1* has been reported to activate NFκB pathway^45, 46^. Taken together, these results suggested an activation in NFκB signaling pathway, indicating a pro-inflammatory state of AST in the early stage of PDm. Besides, previous observation showed increased NFκB level in dopaminergic neurons in post-mortem brains of PD patients and in PD animal models^47, 48^. We suspected that inflammation first emerge in MB_AST and subsequently spread into other cell types, including dopaminergic neurons.

Greatest dysfunctions were observed in MB_OLG (638 DEGs) (Figure 2b). However, the link between OLG and PD is less investigated and remains largely unexplored, which we attributed to the majority of researches were based on post-mortem tissues while lack of attention in PD initiation. Here we were able to depict the transcriptomic changes in MB_OLG early in PD progression. We noticed *Adipor2*, receptor for adiponectin, was upregulated in OLG from PDm (Figure 2d). Deficiency in AdipoR2 can result in inflammation on MG, but in an indirect way^49^. Hence, we assumed that MB_OLG may be the underlying bridge through which *Adipor2* engages in microglial sensitivity *in vivo* toward neuroinflammatory induction. In addition, we observed a large number of NADH dehydrogenase complex assembly related genes were up-regulated in MB_OLG (Table S2). NADH dehydrogenase is a part of the mitochondrial respiratory chain and the main source of intracellular reactive oxygen species (ROS)^50^, whose imbalance causes cell damage via induction of oxidative stress and inflammation.

### Heterogeneity of chromatin accessibility in PDm and WTm brain

To integrate epigenomic data with transcriptomic data, we projected major cell types from snRNA-seq datasets onto sci-ATAC-seq datasets (Methods). Consequently, we successfully projected major cell types onto sci-ATAC-seq datasets. In cerebellum, we identified CB_GC, CB_PC, CB_AST and CB_OLG. In midbrain, we identified MB_DA, MB_GABA, MB_GLU, MB_AST, MB_OLG and MB_MG. In striatum, we identified ST_MSN, ST_GABA, ST_IMN, ST_AST, ST_OLG and ST_MG. In accordant with snRNA-seq guided annotation, we confirmed that promoter and gene body regions of cell type markers were selectively accessible across cell types. ASTs in three brain regions were specifically accessible at *Aqp4* and *Gfap* (Figure 3d, e, f, l, m, n). CB_GCs were accessible at *Ppp2r2c*, with a small subset being specifically accessible at *Pax6* (Figure 2d), indicating an epigenetically distinct subpopulation. CB_PCs shown specific chromatin accessibility in *Car8*, a canonical marker for Purkinje cells (Figure 3l). In midbrain, MB_DAs were accessible in *Th* (Figure 3m). MB_MGs showed distinctly accessibility in *Aif1* and *Serpinf1* while MB_OLGs in *Opalin* and *Mag* (Figure 3m). In striatum, GABAergic neuron markers *Vip* and *Npy* were accessible in ST_GABA and at relatively lower level in ST_MSNs (Figure 3l). Besides, ST_MSNs were also accessible at classic MSN markers: *Tac1* and *Drd2* (Figure 3f). In particular, we observed the accessibility of *Pdgfra* in ST_OPC. To summarize, we provided a valid chromatin accessibility profiles of major cell types in cerebellum, midbrain and striatum at single-cell level.

**Figure 3.**
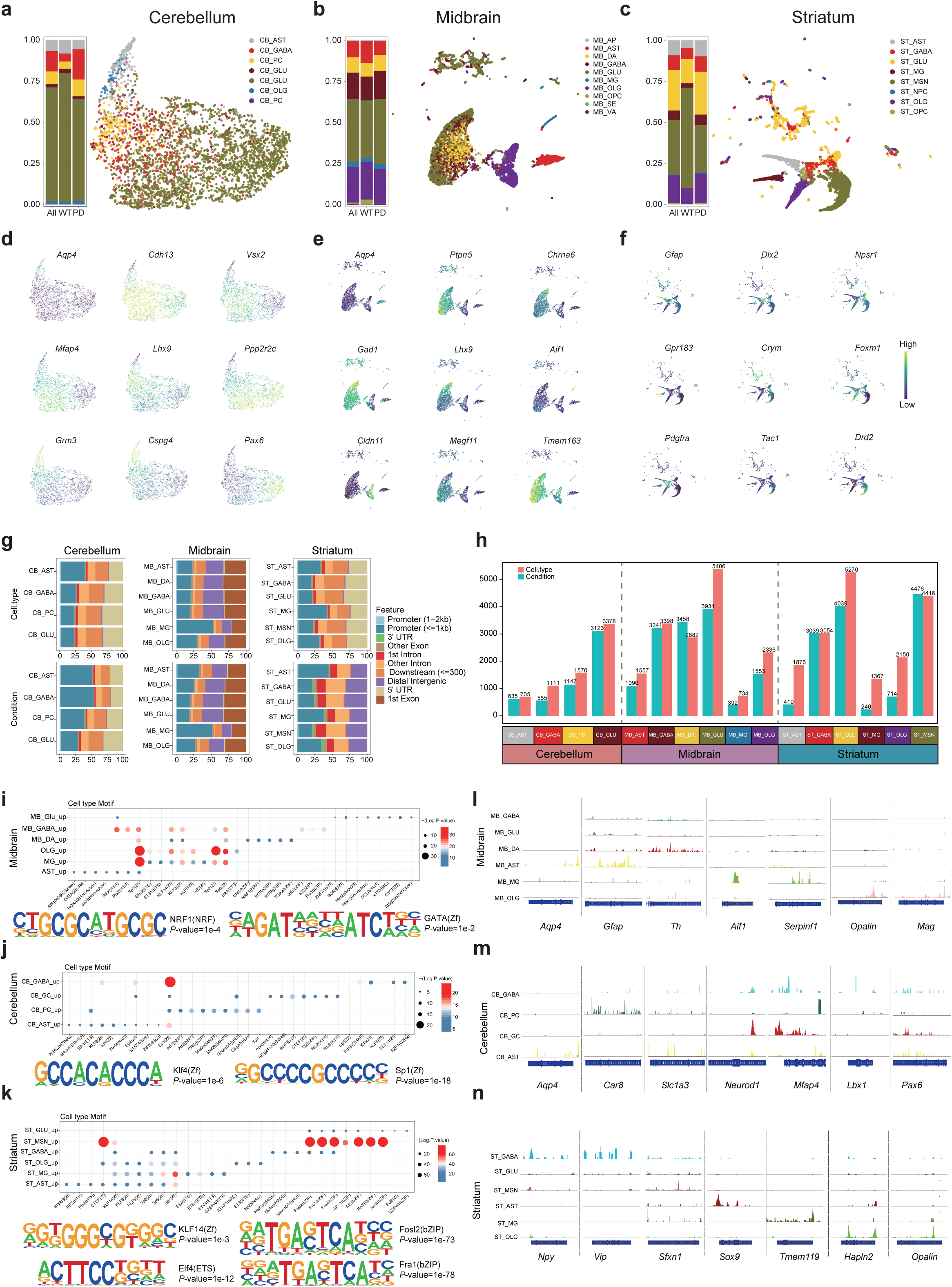
Unsupervised clustering of sci-ATAC-seq datasets of cerebellum (a), midbrain (b) and striatum (c) with cells colored by cell types. Cellular composition is indicated in corresponding barplots on the left of the clustering results. Predicted gene expression profiles using accessibility of reported cell type markers in cerebellum (d), midbrain (e) and striatum (f). Yellow corresponds to high accessibility while purple correspond to low accessibility. g. Genomic distribution of cell type-specific peaks and DARs in PDm at indicated cell types. Left: cerebellum; middle: midbrain; right: striatum. h. The number of cell type-specific peaks and DARs in PDm at indicated cell types. Left: cerebellum; middle: midbrain; right: striatum. Enrichment of transcription factor motifs within cell type-specific peaks at indicated cell types in midbrain (i), cerebellum (j) and striatum (k). Selected transcription factor motifs sequences found in corresponding brain regions are shown. Selected Integrative Genomics Viewer (IGV)^92^ screenshots of representative regions showing the chromatin accessibility of cell type markers in midbrain (l), cerebellum (m) and striatum (n).

We next calculated differentially accessible regions (DARs) for each cell type to further explore their biological functions. We first performed motif enrichment using cell type specific DARs using HOMER^51^. In midbrain, MB_DA-specific DARs showed highly specific enrichment with motifs for CRE, NRF1, RORa, RORg and TGA2, among which *Nrf1* has been confirmed in midbrain dopaminergic neurons to play a neuroprotective role against excessive reactive oxygen species^52^. We noticed GATA3 footprint enrichment (Figure 3i, denoted as GATA(Zf),IR4) in MB_AST DARs, suggesting certain regulations of *Gata3* in MB_AST. In line with our result, a previous study found that GATA3 was able to promote the neurogenic potential in astrocyte^53^. In striatum, DARs of neuronal populations (ST_MSN, ST_GABA and ST_GLU) specifically enriched in motifs for bZIP TF family members: Fosl2, Fra1 and Fra2. Particularly, we noticed that motif of AP-1 were distinctly enriched in ST_MSN DARs (Figure 3k). AP-1 protein complex governs a wide range of cellular processes spanning proliferation, development and apoptosis^54^. AP-1 consists of Fos and Jun, along with activating transcription factor (ATF) ^55^. We found that motifs of Fos, Jun and Atf (Fosl2, Fra1, Fra2, JunB BATF and Atf3) were also enriched in ST_MSN (Figure 3k), further indicating a strong activity of AP-1 in ST_MSN. Striatal glial cells showed relatively stronger enrichment than neurons in Kruppel-like factors family (belongs to Zf family) such as Klf14, Klf3 and Klf5. ETS family members (Elk4, ETS1, ETV4 etc.) were seemingly involved in the normal function of ST_MG. Among them, ETS1 has been reported to be localized with reactive microglia and was ubiquitously expressed in AD brains^56^. In cerebellum, CB_ASTs were likely to be regulated by Klf4 due to the prominent enrichment of corresponding molecular footprint (Figure 3j). *Klf4* has been reported to regulate astroglial and microglial activation^57–59^, further validating our cell type identification.

### Identification of PD biomarkers leveraging multi-omics approach

Genomewide association studies have identified a long list of PD-relevant genomic regions, yet the development of practical treatments for PD remains challenging. Biomarkers with good sensitivity and specificity can aid the diagnosis, progression monitoring and therapies development of PD. Herein we, with both transcriptomic and epigenomic landscape of the pathological brain, sought to identify putative biomarkers at early stage of PD, allowing novel disease interventions to start earlier. Concretely, we aimed to identify changes in both gene expression level and chromatin accessibility level between PDm and WTm brain. In total, we identified three classes of biomarkers for PD (Table S5): bmET (biomarkers that significantly changed at both epigenetic and transcriptomic level), bmE (biomarkers that significantly changed solely at epigenetic level) and bmT (biomarkers that significantly changed solely at transcriptional level), which could be subdivided into two categories each, depending on if that gene was up- or down-regulated at transcriptomic and epigenetic level. We assumed that bmET, bmE and bmT correspond to high-fidelity PD biomarkers, early stage PD biomarkers and candidate PD biomarkers respectively. Of note, we mainly focused on midbrain because PD pathogenesis first occurs in midbrain among three brain regions we investigated.

We first inspected high-fidelity biomarkers (bmET). In MB_DA, *Cox5a, Rcan2, Glul, Atp1b1, Calm1* and *Ctsl* were identified as bmET_up (Figure 4a, b). *Cox5a*, encoding a subunit of terminal enzyme of the mitochondrial respiratory chain, whose up-regulation represents an increased cellular respiration rate and overactivity in electron transport chain^60^, which might be related to ROS production and oxidative stress in PD MB_DA. From an another perspective, we may also be able to interrogate the proteostasis by *Ctsl*, whose product is a lysosomal protease that can degrade α-syn amyloid fibrils^61^. Besides, another bmET *Calm1* encodes a calcium binding protein that regulates a large number of cellular activities. Adding to our speculation, CALM1 has been reported to interact with α-syn and affected its aggregation, suggesting its potent role in PD pathologies^62^. Hence, we proposed that the transcriptomic and epigenetic changes in *Calm1* can be applied for the tracking of deterioration level in PD patients. Specifically, we identified *Scg2* and *Atp1b1* as bmETs in all neuronal cell types in midbrain (MB_GLU, MB_GABA and MB_DA). *Scg2* is engaged in the sorting and docking of neuropeptides into secretory vesicles^63^ and *Atp1b1* encodes a catalytic subunit of Na^+^/K^+^ -ATPase that is responsible for the establishment and maintenance of the membrane potential by regulating Na^+^ and K^+^ flows^64^. In AST, bmET_up (up-regulated bmET) includes *Bcl2*, *Shisa4*, *Dnm3* and *Prex2* (Figure 4a, b; Table S5), among which *Bcl2* and *Shisa4* are involved in NFKB signaling pathway^65^, suggesting their potential to be biomarker for neuroinflammation in PD; Additionally, *Dnm3* plays a part in vesicular transportation, in particular endocytosis^66^, pointing to its association with the ability of AST to maintain extracellular homeostasis. Though no direct association is identified between *Prex2* and PD, *Prex2* has been linked to motor coordination in cerebellum^62^. However, bmET_down (down-regulated bmET) were only identified in MB_GLU (*Dst, Gls, Vegfa, Nav1 etc.*) and MB_GABA (*Unc80*, *Hnrnpu*). In MB_GLU, *Gls* fundamentally regulates glutamate synthesis and its down-regulation, as indicated both transcriptionally and epigenetically, pointed to a possibly defective glutamine level. In MB_GABA (Figure 4c, d), *Unc80* encodes a subunit of a voltage-independent “leak” ion-channel complex, forming and sustaining the resting potential^67^. The down-regulated of *Unc80* in MB_GABA suggests a damaged competence to maintain resting membrane potential.

**Figure 4.**
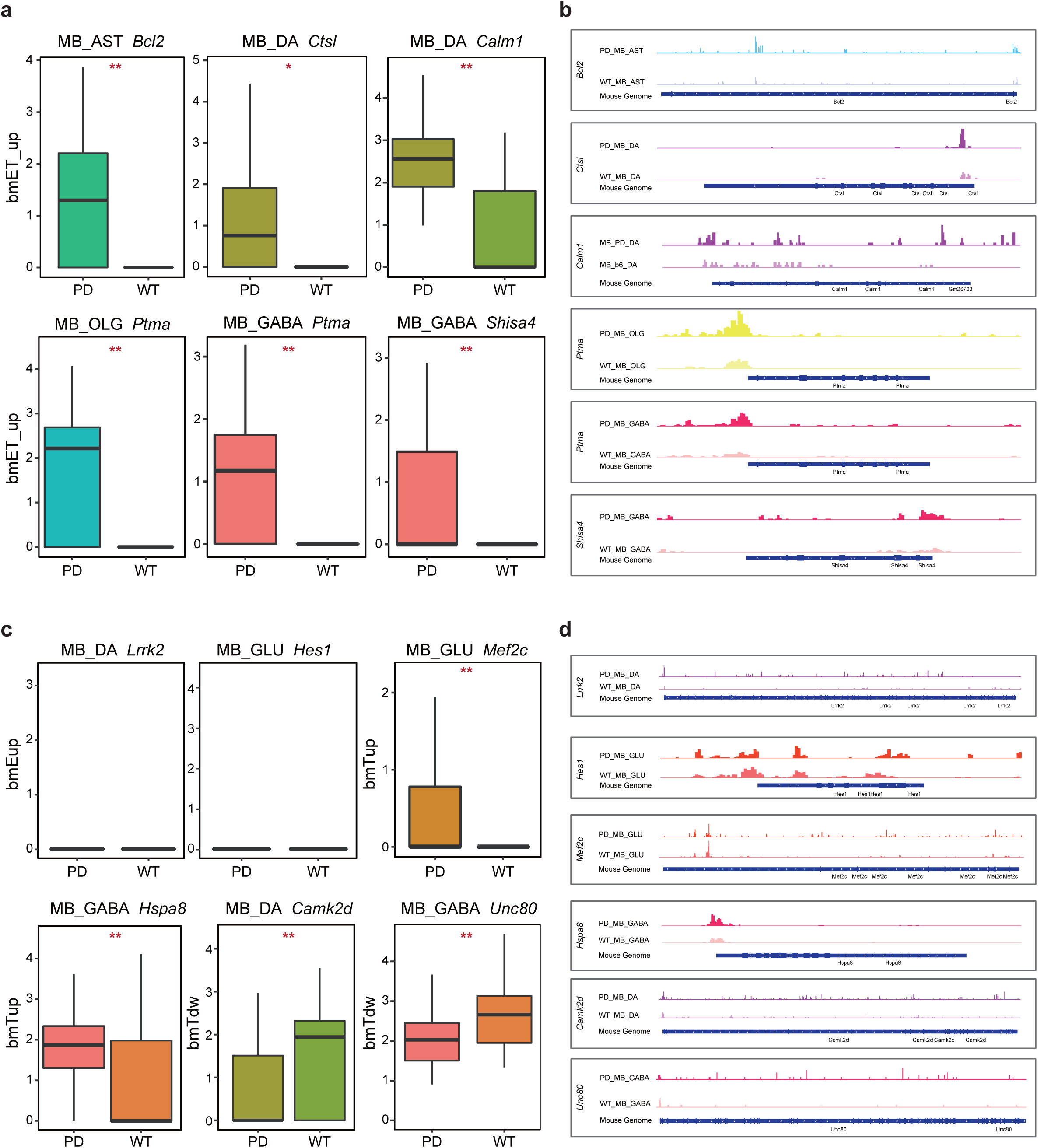
a. Barplot showing the expression pattern of bmET_up genes between PD and WT states in cell types among midbrain. b. Selected IGV^92^ screenshots of representative regions showing the chromatin accessibility of bmET_up genes between PD and WT states in cell types among midbrain. c. Barplot showing the expression pattern of bmT_up, bmE_up, bmT_down, and bmET_down genes between PD and WT states in cell types among midbrain. d. Selected IGV^92^ screenshots of representative regions showing the chromatin accessibility of bmT_up, bmE_up, bmT_down, and bmET_down genes between PD and WT states in cell types among midbrain.

We next identified several bmE_up which are closely associated with nervous system development, including 23 transcription factor encoding genes (Table S5). It is worth noting that *Lrrk2*, a profoundly investigated genes where multiple mutations have been associated with PD, was also identified as bmE_up in MB_DA (Figure 4c, d). A series previous studies provided inconsistent result concerning the relationship between LRRK2 and α-syn, some of which showing LRRK2 deteriorate α-synucleinopathy^62, 68^ while some reporting minimal impact of *Lrrk2* in α-syn aggregation^69, 70^. These observations suggested that complex mechanisms underlie the LRRK2-mediated exacerbation of α-syn neuropathology. Here we captured the up-regulated chromatin accessibility of *Lrrk2* in PD, indicating the epigenetic changes may be an unexplored contributor to α-syn pathologies. We also proposed a list of bmE_down which are closely associated with nervous system development, including six transcription factor encoding genes: *Hes6, Dbx1, Nr2c2, Etv4, Hes1* and *Nfia* (Figure 4c, d, Table S5).

Beside, we also detected a significant proportion of bmT, composing of both bmT_up and bmT_down. In MB_GLU, we identified *Mef2c* to be bmT_up (Figure 4c, d). *Mef2c* encodes a repressor whose expression in excitatory neurons regulates excitatory/inhibitory synapse density principally in a cell-autonomous way^69^. In MB_DA, bmT_up includes *Hspa8* whose product reduces the cellular toxicity as well as intercellular transmission of α-syn fibrils^71^ (Figure 4c, d). The elevated transcriptional expression along with unchanged chromatin accessibility of bmT_up like *Mef2c* and *Hspa8* suggested that they may be regulated by *trans*-acting factors, for example, remote enhancers. Among bmT_down, we found genes that closely associated with nervous system dysfunctions (*Camk2d*, *Il1rapl1, Kcnq2, Nrxn3, Shank1 etc.*). *Camk2d* was a specific bmT_down in MB_DA (Figure 4c, d). *Camk2d* encodes a subunit of calmodulin-dependent protein kinase but has not been linked with PD pathophysiology. *Kcnq2*, encoding a subunit of potassium ion channel whose defect, is a specific bmT_down in MB_GLU.

To summarize, we identified three categories of biomarkers for PD: high-fidelity biomarkers (bmET), early stage biomarkers (bmE) and candidate biomarkers (bmT). High-fidelity biomarkers were supported by both transcriptomic and epigenetic profiles. Early stage biomarkers were exclusively observed at epigenome level. Based on the postulation that chromatin becomes accessible prior to transcription initiation, we proposed the chromatin accessibility of the promoter and gene body regions of these genes can also be indicative of PD. As for candidate markers that solely were supported transcriptionally, we suspected the involvement of trans-regulatory factors.

### Identification of cell types closely associated with PD

We next sought to locate certain cell type that significantly contribute to early PD progression. To achieve this, we performed overrepresentation analysis using PD-risk genes retrieved from DisGeNET ^72^. Most of collected PD-risk genes were identified based on GWAS conducted on PD patients at late stage, thus allowing us to explore the early malfunctions of these genes. Globally, DEGs between PDm and WTm showed significant enrichment scores in the majority of inspected cell types. In midbrain, PDm up-regulated genes in MB_DA showed no enrichment for PD-risk genes, indicating that at early stage of PD, transcriptional dysfunctions in MB_DA showed little consistence with that at late stage. (Figure 5a).

**Figure 5.**
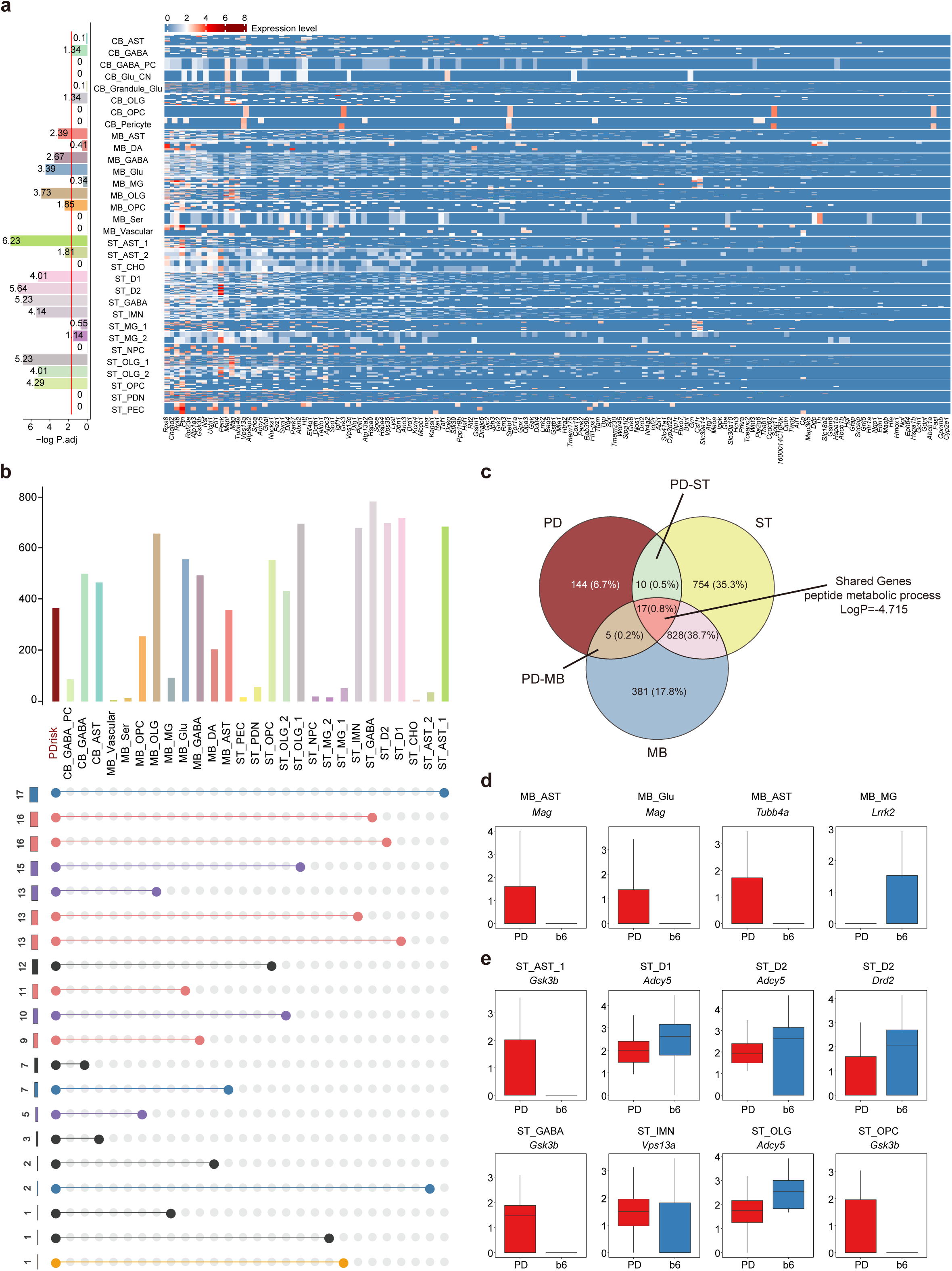
a. Overrepresentation analysis of PD-risk genes. Heatmap: Expression of PD-risk genes in cell types from indicated brain regions. Bar graphs: numbers indicate log-transformed *P*-values, the red line indicates significance (*P*-value=0.05). b. Intersection visualization of PD-risk genes and DEGs in each cell types. Upper bar graphs showing the number of DEGs in indicated cell types. Left bra graphs: the number of element in indicated intersections. c. Overlap of PD-risk genes and PD up-regulated genes in midbrain and striatum d. Expression profiles of selected DEGs in midbrain e. Expression profiles of selected DEGs in striatum

Because AST has been profoundly reported to contribute to PD progression by transferring *α*-syn to dopaminergic neurons, we next inspected the enrichment level of AST and found that PD-risk genes were enriched in midbrain and striatal AST populations (including MB_AST, ST_AST_1, ST_AST_2), but not in cerebellum (Figure 5a). We also noticed that oligodendrocyte lineage (MB_OLG, MB_OPC, ST_OLG_1, ST_OLG_2, ST_OPC and CB_OLG) in midbrain and striatum, holding a previously obscure role in PD pathology, displayed strong enrichments of PD-risk genes to varying extents, indicating putative unexplored clues that lead to early PD initiation and development. Concretely, ST_OLG_1 had a enrichment score of 5.23, followed by 4.29 for ST_OPC, 4.01 for ST_OLG_2, 3.73 for MB_OLG, 1.85 for MB_OPC and 1.34 for CB_OLG (Figure 5a). In conclusion, we observed that most midbrain and striatal cell types were widely perturbed at transcriptomic level. Nonetheless, only 2 cerebellar cell type (CB_GABA, CB_OLG) was enriched for PD-risk genes, which conformed to prior knowledge because cerebellum was not severely affected until Braak stage V^73^.

Next, to evaluate the overlapping between PD-risk genes and DEGs up-regulated in PDm in midbrain and striatum (MB_DEG and ST_DEG), we performed intersection analysis (Figure 5c), resulting in 17 genes (*Snca, Park7, Uchl1, Chchd2, Synj1, Atp6ap2, Rpl23a et al.*) shared by all three datasets (hereafter termed as shared genes), five genes (*Lrrk2, Sod1, Sod2, Mag* and *Tubb4a*) specifically shared by PD-risk genes and MB_DEG (hereafter termed as PD-MBs) while 10 genes (*Drd2, Vps35, Gsk3b, Taldo1* et al.) specifically shared by PD-risk genes and ST_DEG (hereafter termed as PD-STs). Shared genes were mainly involved in peptide metabolic process. In PD-MBs, we noticed *Sod1* and *Sod2* were up-regulated in MB_OLG (Table S2). *Sod1* and *Sod2* are responsible for the elimination of radicals^74^ thus indicating the *α*-syn aggregation may lead to the overproduction of radicals in OLG. *Lrrk2*, whose mutations can disrupt the normal expression of pro- and anti-inflammatory cytokines^75^, was found to be down-regulated in MB_MG (Figure 5d), indicating a dysfunction of the regulation of neuroinflammatory responses in MG. *Tubb4a* was up-regulated in MB_AST (Figure 5d). In PD-STs, we noticed *Vps35* were specifically enriched in ST_AST_1 of PDm (Table S2). VPS35 is a major component of retromer that regulates retrograde transport of proteins from endosomes to the trans-Golgi network. Besides, VPS35 protected mice from neurodegeneration by suppressing *α*-syn expression^76, 77^. In particular, we found that *Drd2*, encoding a dopamine receptor, showed lower expression in ST_MSN_D2 (Figure 5e), suggesting the impaired capability of ST_MSN_D2 of dopamine intake.

## Discussion

We reported 46,146 single-cell transcriptomic and 28,347 epigenetic profiles on human *α*-syn knock-in PD model mouse and matched control. Previous studies mainly focused on specific cell types (dopaminergic neuron, astrocyte or microglia), limiting the investigations on the contributions of intercellular interplays to PD pathogenesis. Here we provided a full picture of both transcriptome and epigenome dysfunctions at early stage of Parkinsonism. In total, we identified major cell types across three brain regions we investigated: GABAergic neurons, glutamatergic neurons, dopaminergic neurons, serotonergic neurons, AST, OLG, OPC, MG, where transcriptionally distinct subpopulations of certain cell types were further identified. For instance, cerebellar Purkinje cells were a subtype of GABAergic neurons. Most of the transcriptomic cell types were epigenetically distinct, suggesting the consistency of our snRNA-seq datasets and sci-ATAC-seq datasets.

In midbrain and striatum, we noticed that DEGs were mainly expressed in neuron axon terminals where the *α*-syn locates and aggregates. Hence, together with the above results, we speculated that at early stage of PD, the translation activity was globally promoted and the splicing events were altered in neurons, jointly triggered enhanced protein degradation pathways. However, a preceding study, by the assessment of post-mortem brain tissue from 70 to 80 years old patients and middle-aged controls using q-PCR, found that *RPL* and *RPS* genes were not detected as significantly changed in SN at Braak stages 1–2 and down-regulated at subsequent stages, which indicated a decline in translational activities^78^. Contradicting to our results, we reasoned that 6-month-old PDm is in an earlier period (before Braak stage 1) of Parkinsonism. The amount of *α*-syn oligomerization may not be sufficient to cause severe deregulation of proteostasis network early in pathological progression of PD and cells are trying to neutralize the disturbing influences of aggregation. Also, bulk measurement in previous study may mask the heterogenous changes within tissue, as a result of which, our results may provide a delineation of the molecular changes in PDm in a finer level.

Membrane trafficking was also affected by oligomerized *α*-syn. Under normal circumstances, peptides for secretory protein will be translated and folded in ER and later enter Golgi apparatus for further modification, followed by secretion in the form of vesicles. The misfolded polypeptides, in our case, misfolded *α*-syn, will be withdrawn from Golgi to ER and undergo subsequent ER-associated degradation (ERAD) or autophagy^79^. In PDm we noticed an intensified retrograde trafficking from Golgi to ER in response to anomalous forms of *α*-syn. Mutant *α*-syn is less susceptible to protein degradation pathway and capable of impairing exocytosis through binding to Rab3a, otherwise hindering the release of neurotransmitters. We also raised a plausible conjecture that the potential Rab26-Rab26os regulatory network and its role in autophagosome genesis.

By interrogating the dysfunctions of channelosome, we noticed impaired calcium homeostasis in ST_MSN and CB_PC. Furthermore, ST_MSN under pathologic condition may has difficulties in the formation of action potential due to descendent Na^+^ influx and anomalous intracellular Ca^+^ signals, suggesting changes in downstream direct and indirect pathway in motor circuits. Importantly, MB_DA showed defective competence of releasing neurotransmitters, which may be another contributing factor of reduced striatal dopamine level.

The initial role of glial cells in PD has long been promoted^69^. Here we observed the transcriptional up-regulation of glutamatergic receptors and transporters in midbrain and/or striatal ASTs, which indicates an excessive accumulation of glutamine in extracellular space or elevated post-synaptic uptake by neurons, potentially leading to hyperactivate phenotypes that demonstrated in early-stage PD^80^. We also noticed the elevated transcriptional expression of NFκB pathway activators (*Bcl2*, *Csnk2a1* and *Chuk*) in PDm MB_AST and MB_MG, based on which we raise the possibility that NFκB signaling pathway was activated even at early stage of PD and this may be the underlying mechanisms regarding neuroinflammatory responses.

Due to the challenge to identify early stage markers for PD to help diagnosis and novel treatments, we identified putative biomarkers, which we divided into three classes: high-fidelity biomarkers, early-stage biomarkers and candidate biomarkers. high-fidelity biomarkers displayed positively-related transcriptional and epigenetic profiles that reciprocally validating each other. Early-stage biomarkers showed the ability to capture PD signals where transcriptomic profiles fail, whereas candidate biomarkers suggested the engagement of trans-regulatory factors that need to be further validated. We also attempted to bridge the gap between genotype and phenotype through overrepresentation analysis of PD-risk genes. Importantly, We noticed that at early stage of PD, the alterations of MB_DA showed little accordance with that found in brain tissue from post-mortem subjects, highlighting the necessity of researches into early stage of PD.

## Materials and methods

### Tissue dissection and nuclear extraction

6-month old PD model mouse (B6;C3-Tg(Prnp-SNCA*A53T 83Vle/J, Jackson Stock No:004479) and recommended control (B6EiC3Sn.BLiAF1/J, Jackson Stock No:003647) were purchased from the Jackson Laboratory. After the animals were sacrificed, brain tissues (including striatum, midbrain and cerebellum) were isolated, quickly froze in liquid nitrogen and stored in liquid nitrogen until library construction. The tissue was taken out from liquid nitrogen and thawed, cut into small piece and transferred into 2ml Dounce Tissue Grinder with 1.5ml 1× tissue homogenization that was comprised of 30mM Cacl2, 18mM Mg(Ac)2, 60mM Tris-HCl (pH 7.8), 320mM sucrose,0.1%NP-40 and 0.1mM EDTA. Tissue was stroked with the loose pestle 15 times and filtered with the 70um cell strainer to remove the cell debris and large clumps, after which tissue homogenate was transferred into the cleaned 2ml Dounce Tissue Grinder again, stroked 15 times with the tight pestle, and then filtered with a 40um cell strainer. The nuclear extraction was spun down in 500g, 4°C for 10min. The supernatant was carefully discarded and the nuclear precipitation was resuspended with PBS containing 0.1% BSA and 20U/ul RNase Inhibitor. Cell count was then performed to calculate the concentration of nuclear suspension.

### Single-nucleus RNA library construction and sequencing

After nucleus extraction, we stained the nucleus with DAPI and counted using microscope to ensure the integration and individuality of the extracted nuclei. Next, we washed the nucleus using PBS containing 0.1%BSA and 0.2U/ul RNA inhibitor at 500g, 4°C for 10min. The nucleus precipitation was resuspended in PBS containing 0.1%BSA and 0.2U/ul RNA inhibitor to a concentration of 1000cells/ul. 16 ul of the nuclei suspension were used to perform 10x RNA library construction following the Single-Cell 3′ Gel Bead and Library V2 Kit guidance. In order to be compatible with BGISEQ-500 sequencing platform, libraries conversion was performed using the MGIEasy Universal Library Conversion Kit (App-A) (Lot: 1000004155, BGI).

### sic-ATAC library construction and sequencing

We performed the sci-ATAC-seq as previously described^81^ with the following adjustments. After the nuclei were extracted, 350,000 nuclei were resuspended using PBS containing 1% BSA and then were equally distributed into 96-well plates. 7 ul nuclei resuspension were added to each well, before which, 2 ul of 5x TAG buffer and 1 ul of Tn5 transposase with different barcodes were already loaded to each well. Thus, the total volume of each well was 10ul. Then transposition were performed in PCR instrument for 30 min at 37°C. Transposition reaction was terminated with the addition of 40 uM EDTA at 10ul/well. Subsequently, the nuclei were mixed together and stained with DAPI, followed by flow sorting. DAPI-positive cells were sorted into 96-well plates with each well containing PCR primers with different indexes. 20-25 cells were sorted into each well. After sorting, the nuclei were briefly centrifuged. Then, 1ul of 0.2% SDS were added and aspirated within each well, followed by incubation at 55 degrees for 7min in the PCR machine. Afterwards, 1ul of 10% Triton-X was added, aspirated and placed at room temperature for 5min to neutralize SDS. Finally, 10ul NEBNext® High-Fidelity 2X PCR Master Mix were added and following the PCR amplification, with 22 cycles. PCR products were purified and performed the library construction. After quality control, the libraries were sequenced on BGISEQ-500 platform.

### Data demultiplexing and quality control

*snRNA-seq:* We first used Cell Ranger 3.0.2 (10X Genomics) to process raw sequencing data and then Seurat^82^ was applied for downstream analysis. Before we started downstream analysis, we focus on four filtering metrics to guarantee the reliability of our data. (1) Genes that are detected in less than three cells were filtered to avoid cellular stochastic events; (2) Cells whose percentage of expressed mitochondrial genes are greater than 10% were removed to rule out apoptotic cells; (3) Cells whose UMI counts are greater than 10000 were removed to filter out the doublet-like cells; (4) Cells whose detected genes are out of the range of 200-4000 were removed.

*sci-ATAC-seq:* SnapATAC^83^ was applied for sci-ATAC-seq dataset processing. Firstly, barcodes whose Hamming distances to the whitelist were less than four were regarded as valid and reads associated with valid barcodes were retained. Secondly, clean fastq files were then aligned to GRCm38.p6 (GCF_000001635.26) using bwa mem. Read entries with MAPQ lower than 30, whose fragment lengths were out of the range of 30-1000 and that were not properly paired were removed. After alignment and filtration, necessary information, including meta data, cell-by-bin count matrix, was used to wrapped up into snap-format files. Next, snap-format files generated from the same brain region (including PDm and WTm) were merged and used as input to SnapATAC for further analysis. To ensure the validity of the dataset, we established the following quality control metrics: (1) Cells with fragments and unique fragments greater than 10000 and 800 respectively were retained; (2) Cells whose fragments in promoter ratio (FiPR) were in the range of 0.2-0.8 were retained. Promoter regions were defined as the 2kb upstream and 10bp downstream of transcription start sites (TSSs); (3) Bins located in blacklist regions (identified by ENCODE^84^), unwanted chromosomes (X, Y, mitochondrion) were removed; (4) Bins whose coverage were in the range of 0 and 95-quantile were retained. Bin coverage was defined as the number of cells that has fragment(s) in a certain bin.

### Clustering

After quality control, we performed clustering in a region-dependent fashion.

*snRNA-seq:* Unsupervised clustering were performed using Seurat v3^85^. A series of pre-processing procedures were performed separately on each sequencing library before clustering. Firstly, normalization was performed employing “LogNormalize”. Concretely, in each cell, raw UMI counts for each gene were divided by the total expression, multiplied by 10,000, and transformed to log space. Next we calculated variance scores for each gene based on dispersion and average expression. The top 2000 genes were defined as highly variable genes (HVGs). Then we applied “FindIntegrationAnchors” and “IntegrateData” functions to integrate all sequencing libraries (including PDm and WTm), followed by the regression of technical noise. Principal component analysis (PCA) was performed using HVGs and principal components (PCs) significance was calculated using the “JackStraw” function. In this case we chose top 20 significant PCs for downstream cluster identification and visualization. Clusters were defined based on k-Nearest Neighbor (KNN) algorithm with k=20. UMAP were used for visualization. For sub-clustering, we followed the pipeline described above but only using chosen cells.

*sci-ATAC-seq:* Diffusion map were utilized for dimensionality reduction. By intuitively observing the pairwise correlation plot of adjacent eigenvectors, we were able to determine the top six dimensions to include for downstream analysis. We construct a KNN graph where k=100, based on which clusters were defined. UMAP were run on the chosen components for visualization.

### Cell type annotation and sub-clustering

*snRNA-seq:* Cell type were assigned by the expression of known cell-type markers retrieved from published researches.

*sci-ATAC-seq:* We applied an snRNA-based method to identified cell identities in sci-ATAC-seq dataset. Briefly, we first quantified gene activity based on accessibility matrix. Gene activity is defined as the number of fragments in bins overlapping with gene body region; Secondly, cell-to-cell counterparts (termed “anchors”) between snRNA-seq and sci-ATAC-seq datasets were calculated using “FindTransferAnchors” in Seurat^85^, which were utilized for the transfer of cell type labels in snRNA-seq dataset to the sci-ATAC-seq dataset. Cells of prediction scores lower than 0.5 were removed.

### Differentially analysis and functional enrichment analysis

*snRNA-seq:* Differentially expressed genes (DEGs) were identified using “FindAllMarkers” function implemented in Seurat^86^. Wilcoxon rank sum test was applied. Gene with adjusted *P*-value (Bonferroni method) lower than 0.05 was defined as DEGs.

*sci-ATAC-seq:* To identified differentially accessible regions (DARs), (1) We first performed peak calling separately for each cell type using MACS2^87^ with options “-f BEDPE -B --SPMR --call-summits”. Peaks with log(*Q-*value) less than four were retained and merged as a standard peakset. Of note, we skipped this step for cell type with cell number below 100 to obtain more robust outcome. (2) “differentialGeneTest” function implemented in Monocle^88^ were used with minor modifications regarding background cells selection. Here for each cluster, we used their neighboring cells in the calculated diffusion space as background signals. For situation that the inspected cluster accounts for more than half of the total cells, the remaining cells will be used as background. (3) ChIPseeker^89^ were used for DARs genomics features and peak-to-gene annotation.

Functional enrichment analysis were performed using clusterProfiler R package^90^ and the BH method was used for multiple test correction. GO terms with a *P*-value less than 0.01 and KEGG term with a *P*-value less than 0.05 were considered as significantly enriched.

### Motif enrichment analysis

HOMER^51^ was used for motif enrichment with options “-len 10 -size 300 -S 2 -p 5 -fdr 5 - nomotif”.

### Overrepresentation analysis

We first retrieved PD-risk genes from DisGeNET^72^ and then performed the hypergeometric test (“phyper” function in R) using the PD-risk genes and DEGs between PDm and WTm in each cell type. 0.05 of *P*-value was used as a threshold to define the significance. Of note, only genes with Score_gda>0.1 were retained and human gene symbols were converted to mouse gene symbols using biomaRt package^91^.

## Data Availability

The data that support the findings of this study have been deposited in the CNSA (https://db.cngb.org/cnsa/) of CNGBdb with accession code CNP0000892.

## Acknowledgement

Dongsheng Chen is supported by China Postdoctoral Science Foundation (grant number 2017M622795). We thank the support of National Natural Science Foundation of China (No. 31702074 and No. 31872309). We are also thankful to the production team of China National GeneBank, Shenzhen, China.

## Authors Contributions

S.P.T., S.D.Z., D.S.C. conceived the project and revised the manuscript; J.X.Z., J.C.Z. W.Y.W. performed data analysis and wrote the manuscript; X.M.L., C.C.C., L.C.L. participated in experiments; F.Y.W. and L.H.L. contributed to data visualization. J.K.L., F.C., Z. H. and X.X. participated in project discussion.

---

**Figure S1.**
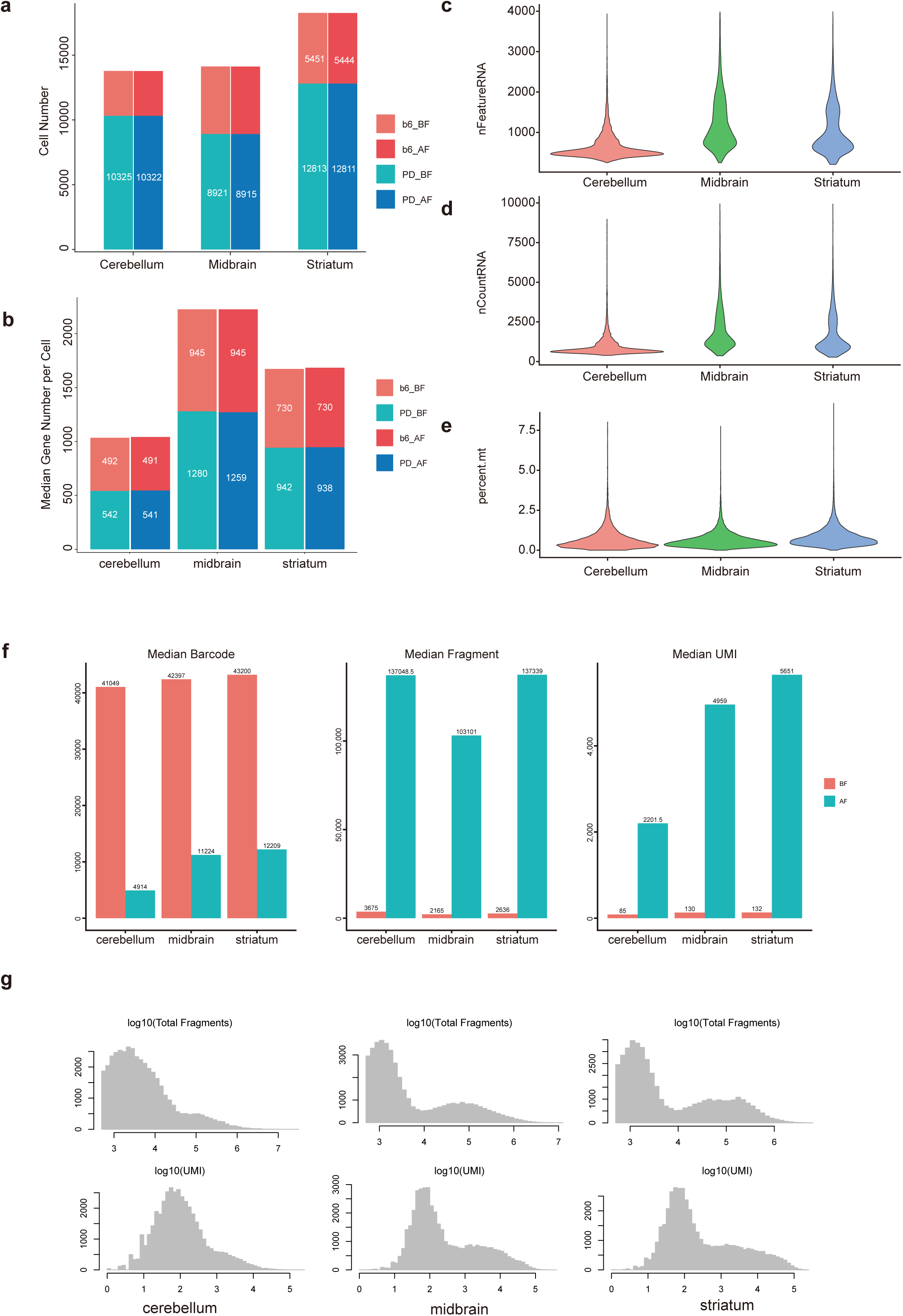
a. The number of cells in snRNA-seq dataset of each brain region before (denoted as BF) and after (denoted as AF) filtering. b. The median of detected genes per cell in snRNA-seq dataset of each brain region before (denoted as BF) and after (denoted as AF) filtering. c. Violin plot showing the number of detected genes in snRNA-seq dataset of each brain region after filtering. d. Violin plot showing the number of UMI in snRNA-seq dataset of each brain region after filtering. e. Violin plot showing the percentage of mitochondrial genes in snRNA-seq dataset of each brain region after filtering. f. The number of cells (left), the median of sequenced fragments (middle) and unique sequenced fragments (right) in sci-ATAC-seq dataset of each brain region before (denoted as BF) and after (denoted as AF) filtering. g. Log-transformed distribution of total fragments (upper panel) and unique fragments (bottom panel) in each brain region. Left: cerebellum; middle: midbrain; right: striatum.

**Figure S2.**
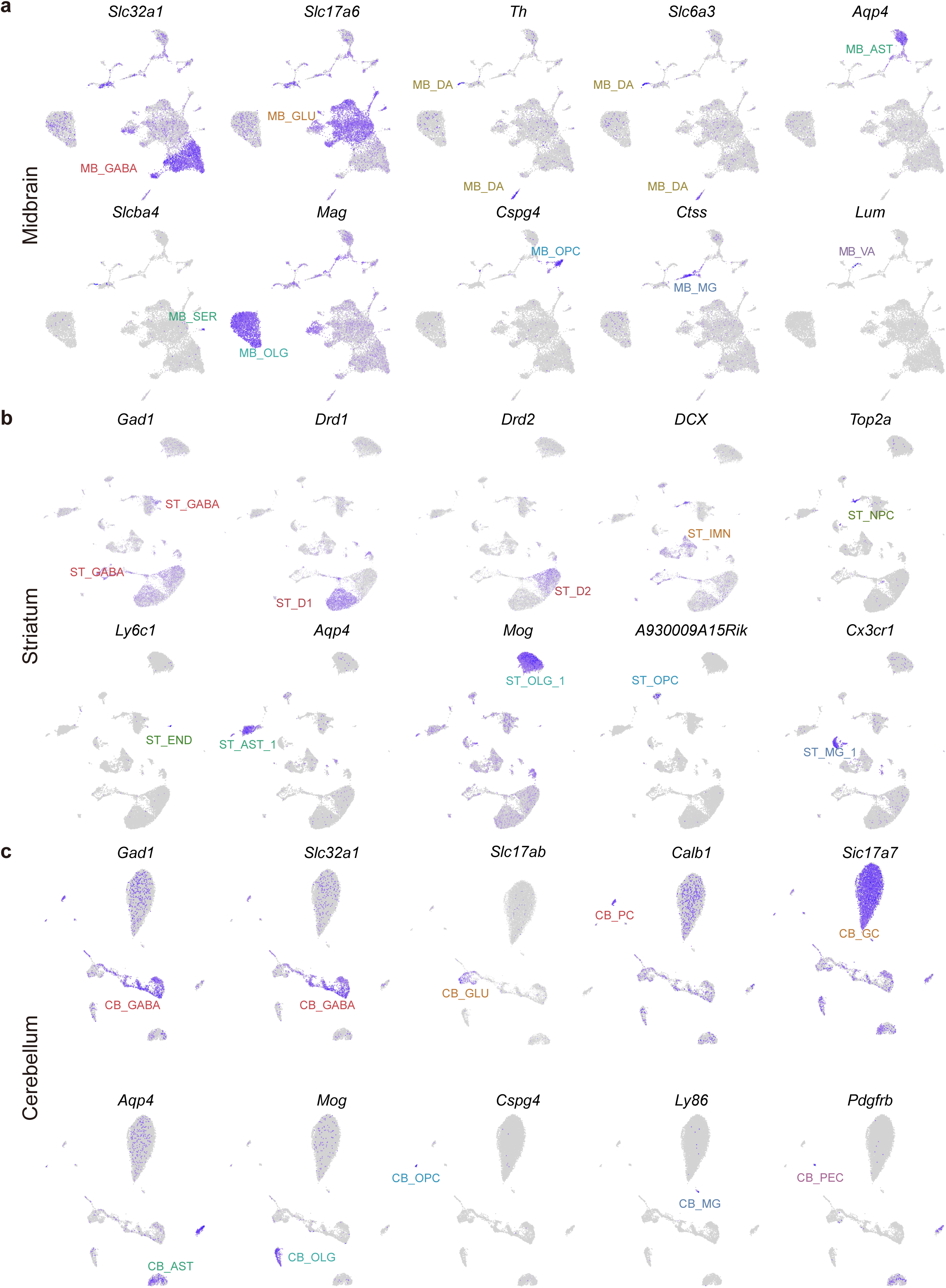
Expression patterns of canonical cell type markers in midbrain (a), striatum (b) and cerebellum (c). Expression level is indicated by shades of blue.

**Figure S3.**
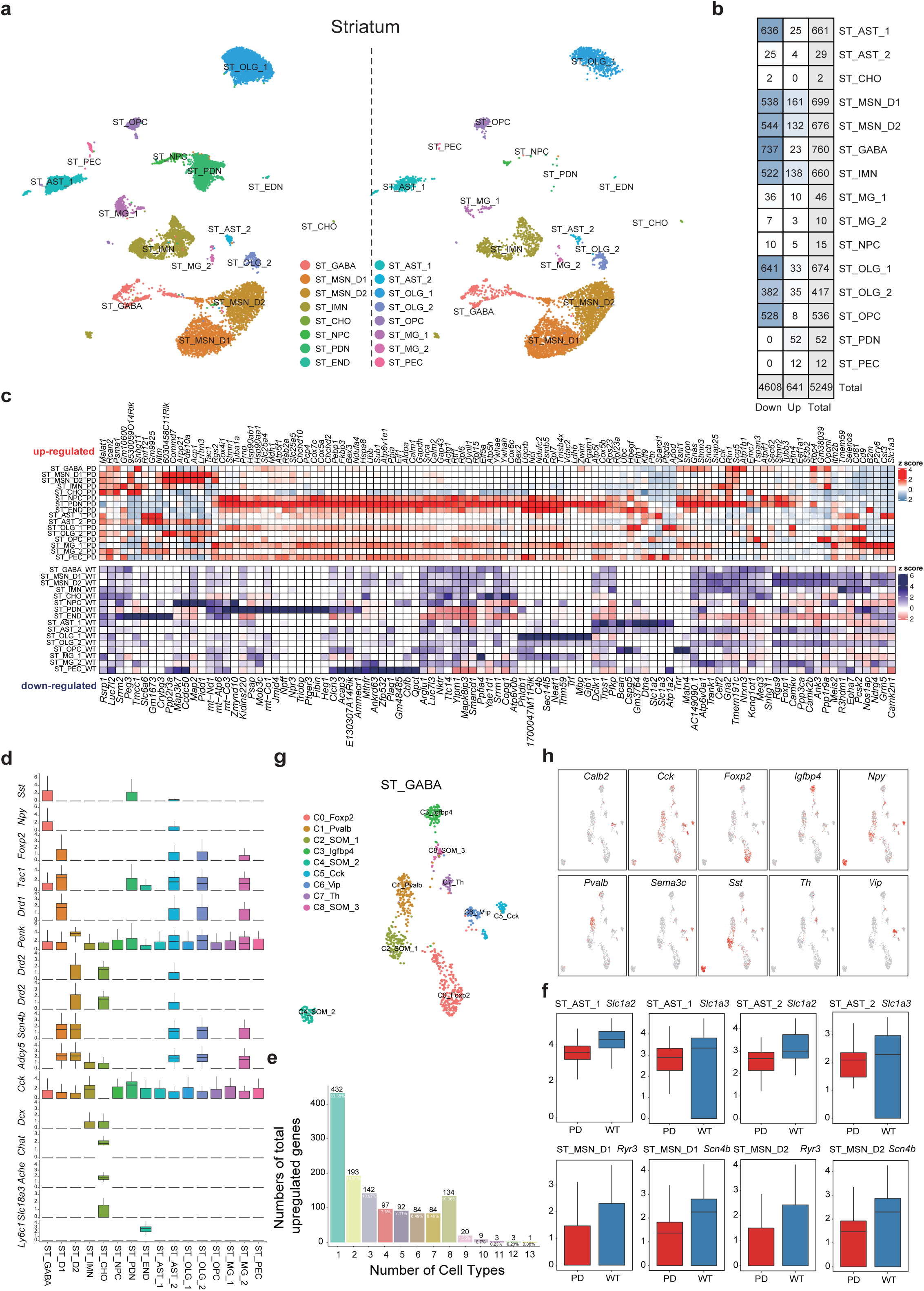
a. Unsupervised clustering of snRNA-seq datasets of striatum, with PD dataset shown in left panel WT dataset shown in the right. Cells are colored by cell type. b. The number of cell type-specific genes c. Expression profiles of cell type-specific genes. d. Expression profiles of selected DEGs e. Sub-clustering of GABAergic interneurons (denoted as GABA in Figure S3a). f. Expression patterns of cluster-specific genes in Figure S3e. g. The numbers of total upregulated genes (y axis) as a function of the total number of cell types in which the upregulation occurs. h. Boxplots showing PD up-regulated genes in indicated cell types.

**Figure S4.**
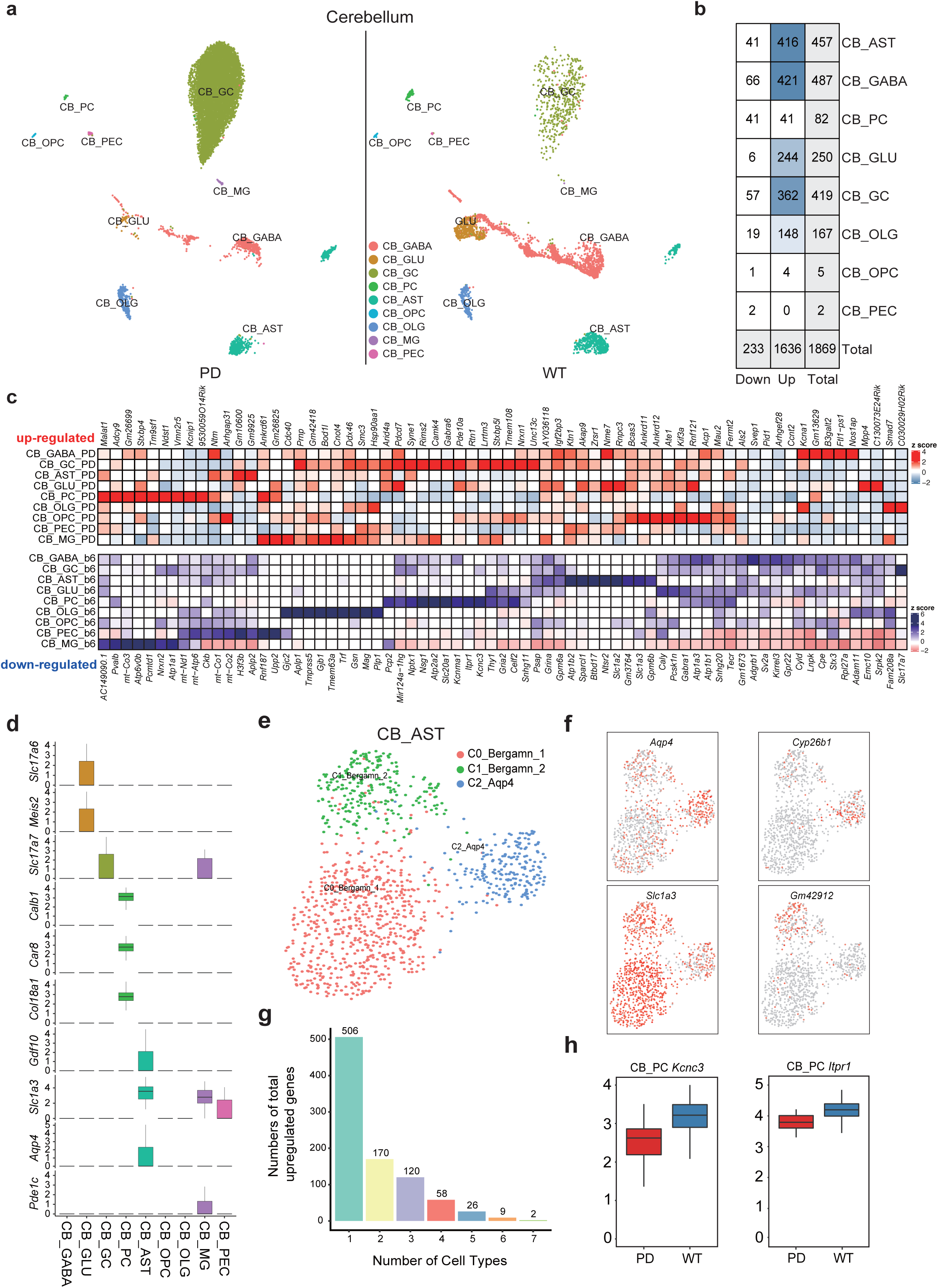
a. Unsupervised clustering of snRNA-seq datasets of cerebellum, with PD dataset shown in left panel WT dataset shown in the right. Cells are colored by cell type. b. The number of cell type-specific genes c. Expression profiles of cell type-specific genes. d. Expression profiles of selected DEGs e. Sub-clustering of cerebellar astrocytes (denoted as AST in Figure S4a). f. Expression patterns of cluster-specific genes in Figure S4e. g. The numbers of total upregulated genes (y axis) as a function of the total number of cell types in which the upregulation occurs. h. Selected GO terms enriched in genes with over three times of occurrence across different cell types.

Table S1. Sample sequencing quality and data quality control information of data generating from snRNA-seq and sci-ATAC-seq.

Table S2. DEGs of clusters in brain regions and DEGs of states between PD mouse and wild type mouse within clusters.

Table S3. DARs of clusters in brain regions and DARs of states between PD mouse and wild type mouse within clusters.

Table S4. Motifs of clusters in brain regions and Motifs of states between PD mouse and wild type mouse within clusters.

Table S5. Biomarkers.

